# Increase of tropical seabirds (*Sula*) in the California Current Ecosystem with warmer ocean conditions

**DOI:** 10.1101/2024.10.14.618240

**Authors:** Tamara M. Russell, David M. Pereksta, James R. Tietz, Maria Vernet, Jaime Jahncke, Lisa T. Ballance

## Abstract

Climate change is impacting marine ecosystems in the California Current Ecosystem (CCE) through physical changes (e.g., an increase in the frequency and intensity of marine heatwaves) that manifest biologically at all trophic levels. We investigated range expansion into the CCE and correlations with environment for a group of tropical/sub-tropical seabirds. We assessed changes in the abundance (2002-2022) of five species from the genus, *Sula* (Cocos, Blue-footed, Red-footed, Masked, and Nazca Boobies), using a novel compilation of four data sources (at-sea surveys, n=82 observations, records from Southeast Farallon Island, n=600, the California Rare Birds Committee, n=593, and *eBird* -a citizen science mobile application, n= 20,529), and looked for relationships with the environment, including broad temporal and spatial scale dynamics (El Niño Southern Oscillation), local conditions where the bird was reported, and potential source conditions from Baja California Sur and the Gulf of California. All five species increased in abundance, and all, with the exception of Blue-footed, exhibited a northward range expansion by as much as 6.8 degrees latitude and increased range area of between 235-1013% within the CCE. There was increased abundance during warmer conditions (El Niño and warm SST) for Cocos, Red-footed, Masked, and Nazca Boobies, and all species increased by 692-3015% after the extreme marine heatwaves that began in late 2013. Our results document a tropical shift in seabirds of the CCE, which may present future challenges to resident species. As marine heatwaves are projected to increase in frequency and intensity, in addition to long-term warming, we hypothesize that these species will continue to expand their range northward in the CCE.

## INTRODUCTION

Species distributions are changing rapidly in response to anthropogenic climate change, due to direct and indirect impacts, including changes in species interactions, prey availability/quality, reproductive habitat availability/quality, and/or physiological restraints (Pinsky et al., 2013; Wingfield et al. 2015; Pinsky et al., 2020). Species moving into or out of an ecosystem may have impacts on the ecosystem structure, such as food web efficiency, stability through diversity, and interspecies competition (Albouy et al., 2014; Bartley et al., 2019; Tekwa et al., 2022).

Seabirds are conspicuous marine predators, and their distribution and abundance patterns can provide information on the underlying marine food web (Fauchald 2009). Changes in seabird distributions can signal perturbations in an ecosystem and new species may become competitors for resources with resident species. Understanding drivers of changing species distributions allows us to better predict how an ecosystem may continue to change with a warming climate and how these shifts may impact existing species in these regions.

The California Current Ecosystem (CCE) is a productive eastern boundary upwelling region (Figure 1). Within the southern sector of the CCE, the Southern California Bight (SCB) is a unique transition zone between cool and warm water ecosystems. The headland of Point Conception, California, is where cool water from the north meets warm subtropical waters from the south (Horn & Allen 1978, Hunt et al., 1980, Hendershott & Winant 1996), and corresponds to the northern or southern extent of many species’ ranges (Hayward & Venrick, 1998, Pitz et al., 2020, Sydeman et al., 2009, Horn & Allen 1978, Briggs & Bowen 2011).

**Figure 1:**
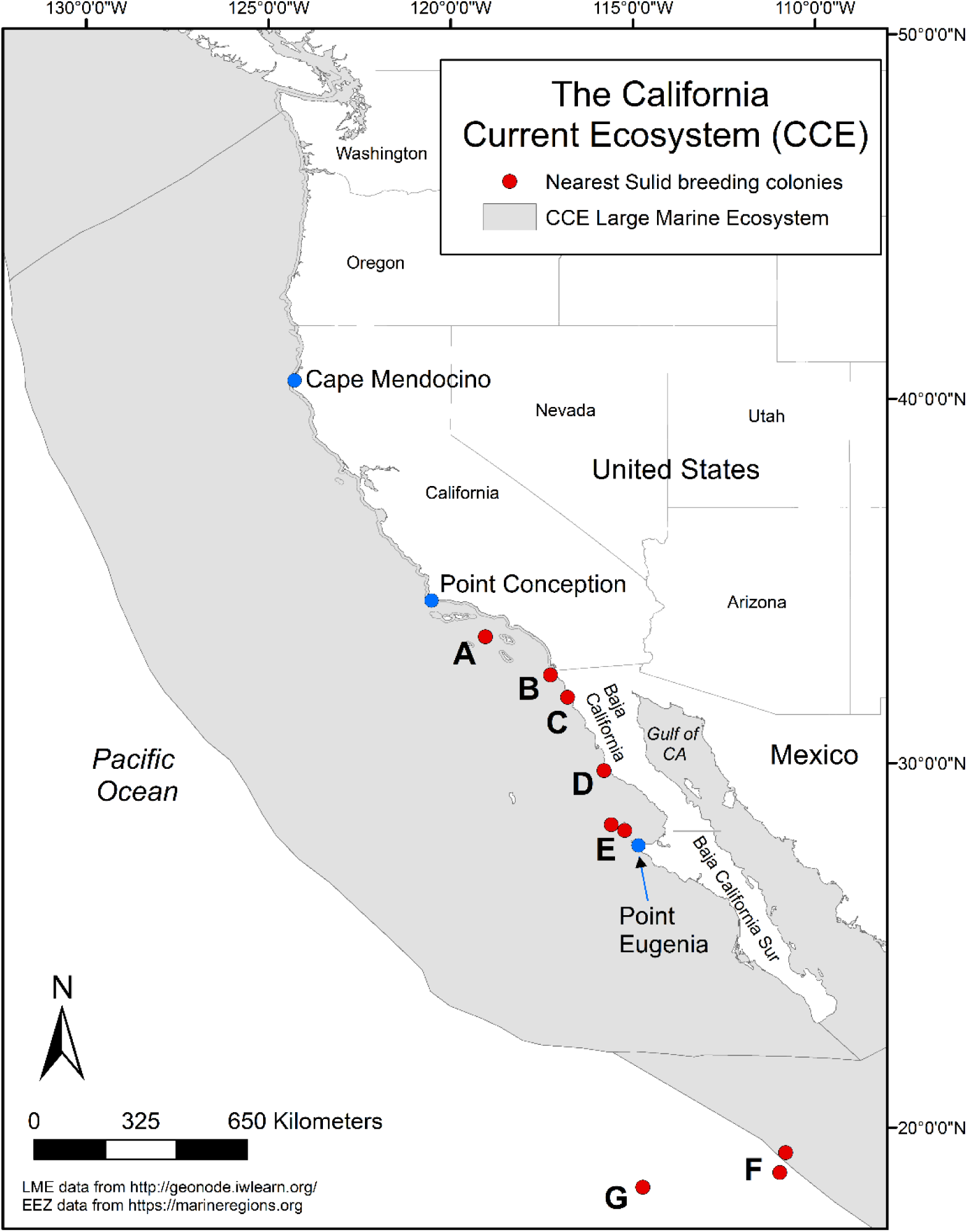
Map of the California Current Ecosystem. The north and central biogeographic regions are separated by Cape Mendocino, the central and south by Point Conception and the south extends down to Point Eugenia. The locations of Sulid breeding colonies in the Pacific are indicated in red and are as follows: (A) Sutil Rock, Santa Barbara Island, (B) Middle Rock, Coronado Islands, (C) Todos Santos Islands, (D) San Jerónimo Island, (E) San Benito and Cedros Islands, (F) San Benedicto and Socorro Islands (inner Revillagigedo Islands), and (G) Clarion Island (outer Revillagigedo Islands).

Climate change is impacting marine ecosystems in the CCE through physical changes (e.g., an increase in the frequency and intensity of marine heatwaves; MHW) that manifest themselves biologically at all trophic levels (Jacox et al., 2020). As ocean temperatures have warmed within the SCB, the cool-warm gradient has blurred, and subtropical species from warmer waters have moved north (Rasmussen et al., 2020, Sagarin et al., 1999, Ainley 1980, Hyrenbach & Veit 2003). For example, elegant terns that typically breed only within the Gulf of California, Mexico, have been shifting north to the southern California, USA, coast to breed during warm years and poor food availability in their usual habitat (Velarde et al. 2015; Veit et al., 2021). In addition, Cocos Boobies (*Sula brewsteri*; previously Brown Booby, *S. leucogaster brewsteri*) and one pair of Blue-footed Boobies (*S. nebouxii*) have recently (2017 and 2022, respectively) expanded their breeding range to Sutil Rock off Santa Barbara Island, California (Howard et al., 2024; Figure 1). Preceding this breeding range expansion, both Cocos and Blue-footed Boobies had expanded their breeding to locations off of the west coast of northern Baja California, Mexico; Cocos to Middle Rock, Islas Los Coronados in 2005 (Whitworth et al., 2007), and Blue-footed to San Jerónimo Island in 2016 (Bedolla-Guzman et al. 2019). Although these breeding expansions are well-documented, the drivers of these expansions have yet to be explained. In addition, sightings of all five Sulids that occur off the Pacific Coast of Mexico are increasing in the CCE, which includes Cocos, Blue-footed, Red-footed (*S. sula*), Masked (*S. dactylatra*), and Nazca Boobies (*S. granti*).

Species within the genus *Sula* are large (wingspans between 140-165 cm), plunge-diving seabirds that are found in tropical and subtropical habitats (Nelson 1978). Due to their size and flight-style, they have high energetic needs and require large amounts of food to sustain them (Ballance 1995; Nelson 1978). Two of these species have pantropical distributions (Red-footed, and Masked), while Cocos, Nazca, and Blue-footed Boobies occur only in the Eastern Tropical Pacific (Nelson 1978). Prior to the expansion and colonization of Middle Rock, San Jerónimo Island, and Sutil Rock, the nearest Sulid colonies were located off Baja California Sur, central Mexico, and within the Gulf of California (Nelson 1978, Whitworth et al., 2007; Howard et al., 2024). As these species mostly live in tropical, lower productivity regions, the impacts of warming temperatures may have disproportionate effects on already low stocks of potential prey, and they may be experiencing competitive pressures to venture farther for resources.

Here, we document the northward range expansion of five Sulids and test our hypothesis that their occurrence in the CCE relates to warmer ocean conditions. Data on ‘rare’ or infrequent species are inherently lacking, therefore including all available data may increase the ability to detect abundance and distribution trends. In this study, we take an integrative approach and utilize structured, semi-structured, and unstructured data to describe changes in Sulid abundance and evaluate their relationships with the environment to better understand the impacts of warming in this upwelling ecosystem.

## METHODS

### Study Area

The CCE is a productive eastern boundary current that stretches roughly 2,800 km from the North Pacific Current (∼50°N) down to Baja California, Mexico (27°N; Figure 1; Checkley and Barth, 2009). Within the CCE are three biogeographic regions –North, Central, South– that have distinct upwelling patterns and biological communities (Ainley, 1976; Ainley et al., 2015; Schipper et al., 2016, Russell et al., 2023). The North (up to 50°N) and Central regions are separated by Cape Mendocino (40.438°N), the Central and South by Point Conception (34.448°N), and the South extends to Point Eugenia (27.848°N, -115.083°W; Checkley and Barth, 2009).

### Seabird data

All data processing, compilation, and analyses were conducted in the R environment (version 4.0.3, R Core Team 2020). Sulids in the CCE are considered vagrants, or species that are rarely seen within a region due to it being outside of its typical range (Stake 2012; Taylor et al., 1994; Mason et al., 2007). As data on vagrants are inherently rare, we used multiple data sources to investigate trends in Sulid abundance and potential environmental drivers (Figure S1 & S2). We utilized structured (at-sea strip transects), semi-structured (summarized surveys and incidental sightings on Southeast Farallon Island; SEFI & *eBird* mobile application; Sullivan et al. 2014), and unstructured data (Rare Bird Committee reports). Semi-structured data include information about the observation, such as duration and distance travelled during data collection, whether the observer recorded data on all species, and other information that provides metadata along with species observations. The integration of public-collected (i.e., ‘citizen science’), semi-structured data with structured survey data has been shown to improve estimates of abundance (Schindler et al., 2020) and population growth rates (Walker & Taylor 2017; Horns et al., 2018; Robinson et al., 2018). We then filled in potential gaps of observations with unstructured data from the California, Oregon, and Washington Rare Birds Committee reports. For each data type, we compiled data on all five booby species reported in the CCE, hereafter referred to as Cocos, Blue-footed, Red-footed, Masked, and Nazca.

#### a. At-sea data

Within the CCE, there are a wealth of data on the at-sea distributions of seabirds that have been collected through temporary or long-term systematic surveys over several decades. These data are collected using standard strip transect methods from ships and aerial platforms (Tasker et al., 1984; Ballance, 2007; Mason et al. 2007) and can be used to estimate density of seabird species within the surveyed transect (Henkel et al., 2007). Data from 21 of these programs were previously compiled into a common format (Leirness et al. 2021; 1980-2017), to which we added additional years from California Cooperative Oceanic Fisheries Investigations (CalCOFI), Applied California Current Ecosystem Studies (ACCESS), and NOAA’s Southwest Fisheries Science Center’s California Current Ecosystem Survey (CCES) resulting in a dataset that spanned 43 years (1980-2022; Table 1S, Figure S1.A). Observations were subdivided into smaller transect segments (mean = 3.91 km) to calculate area surveyed (transect length x transect width; km^2^) and assigned coordinates of each transect segment midpoint.

#### b. Southeast Farallon Island data

There has been avian monitoring at SEFI, California since 1967, including documented daily sightings of bird species. These data include the species, number of individuals, and date of first arrival in any given year, and were summarized as monthly sightings spanning 56 years (1967-2022; Figure S1.B).

During the fall (mid-August to early December), researchers conducted daily counts for Brown Pelicans and Sulids from the lighthouse from 0700 to 0900 when no part of the island was obscured (e.g., fog, clouds, etc.). During other seasons (December to mid-August), the counts were conducted opportunistically. Throughout the year, a 5-minute sea watch was conducted before 0900 with a 30x spotting scope from the houses facing southwest. Additionally, during the fall, a 30-minute sea watch was conducted from the east side of the island late in the afternoon and between 2005-2022, area searches were conducted once in the morning and once in the afternoon, until mid-November when the afternoon survey was dropped. Area searches consisted of a single person walking all accessible parts of the island to search for land birds, but non-land bird species such as Sulids were also recorded.

Dates of arrival and counts were estimated for each unique individual, their identification based on age, sex, and plumage, within a one-month period. For Cocos, an unknown individual (no age or sex information recorded) was assumed to be the same bird as a previously seen Cocos within a 30-day window. For Blue-footed, four juveniles showed up at SEFI in 2013 when there was a large influx of juveniles into California (Rottenborn et al., 2016). Since there were very few Blue-footed reported in the CCE afterwards, the two that showed up as second-year birds in 2014, and returned as adults to the same location each year (2015-2019) were considered two of the four juveniles from 2013. For Red-footed, Masked, and Nazca, individual records were based on California Rare Birds Committee decisions.

#### c. Rare Birds Committee

We used data from three Rare Bird Committees (RBC), volunteer organizations in the U.S. that maintain rare bird records. The public submits records of rare birds (a classification that ranges from 4 occurrences per year in CA, 20 or fewer records during the previous 10-year period in WA, and varies in OR) to the committee and the committee evaluates, confirms (or declines to confirm) observations, and manages an open access records database. We compiled data from the California (CRBC), Oregon (ORBC), and Washington RBC (WRBC) rare bird reports to fill in Sulid occurrences that were not recorded in other data types (https://californiabirds.org; https://oregonbirding.org; https://wos.org; Benson et al., 2022). We only used accepted and verified reports in our analyses. From documentation, the committee also assesses whether reports are new or recurring individuals, and if recurring, they record the date of their first arrival and date last reported. We used the date of first arrival in all analyses. The RBC records used in this study included 576 Sulid observations (CRBC n=504, ORBC n=40, WRBC n=32) spanning from 1935 to 2023 (Figure S1.C). By 2021, all five Sulids (Cocos =2007, Blue-footed =2014, Red-footed & Nazca=2019, Masked=2021) were removed from the CRBC rare bird list. However, all Sulids except for Cocos (removed in 2023 from ORBC and 2018 in WRBC) are still tracked by ORBC and WRBC.

#### d. eBird data

We used citizen science data from the mobile application, *eBird*, which is hosted by Cornell Lab of Ornithology (Sullivan et al., 2014). Citizen science data covers larger spatiotemporal scales than what is economically possible for structured data programs (Conrad and Hilchey 2011; Walker and Taylor, 2017). The use of *eBird* has been growing globally at exponential rates since it was launched in 2002 (Walker and Taylor 2017). These data are publicly available for download (www.ebird.org/science/download-ebird-dataproducts) and can be processed using the ‘auk’ R package (Strimas-Mackey, Miller, & Hochachka, 2017). On *eBird*, an observer can note method of data collection (e.g., stationary, travelling, incidental, or historic records), survey effort (e.g., time and distance travelled), and whether they recorded all species they could identify or were casually noting birds. Once bird lists are submitted, they are flagged if there are anomalous observations (e.g., species not regularly seen in that area or unusually high counts), and regional volunteer reviewers contact the observer for more details and either accept or reject the observation.

We downloaded all *eBird* data for the five Sulids in our study, their hybrids, and Sulid groups (i.e., those that could not be identified to species). This resulted in 204,577 individual records, spanning across all months from 1800 to 2023, which included many historical records (pre-2002 when *eBird* was established).

We downloaded sampling effort data (information on all lists submitted regardless if a Sulid was seen) and connected Sulid observation data to the sampling effort by each unique survey location. We only used *eBird* data that had been reviewed and accepted by regional *eBird* reviewers. We changed each ‘presence-only’ record to one individual, which conservatively estimates species abundance. We calculated the number of lists submitted per day at each location and standardized Sulid observations with this number. This controlled for multiple reports on the same individual(s) submitted at the same location on the same day (Figure S2).

For all four data types, we assigned observations to regions outside and within the CCE. Regions outside of the CCE were not used in analyses but were plotted alongside CCE (Figure S2). We assigned data to the CCE (CCE Large Marine Ecosystem; www.geonode.iwlearn.org) and to the Gulf of California (ww.marineregions.org) using shapefiles with a 20-km buffer to capture coastal observations. We classified remaining data as the “Salton Sea” if they were located within California and south of 34°N, “Inland” if the location was within the remaining parts of California, Oregon, or Washington or anywhere within Arizona, Nevada, and Utah, and “north of CCE” for all northern observations. We also classified data by latitude and biogeographic region-Baja California Sur (hereafter S. Baja) and Southern CCE separated by Point Eugenia, Southern CCE and Central CCE by Point Conception, and Central CCE and Northern CCE by Cape Mendocino (Figure 1).

### Environmental data

To determine potential correlates of Sulid abundance, we used a suite of environmental variables and indices to represent the broader region, the CCE, and potential source locations (conditions in S. Baja and Gulf of California) including the Oceanic Niño Index (ONI), sea surface temperature (SST), Bakun Upwelling Index, and air temperature.

For broad-scale (3-4 year) conditions in the North Pacific, we used ONI as a measure of the El Niño Southern Oscillation (ENSO). ONI is calculated by a three-month running average of SST, where high values (>0.5) are warmer, El Niño conditions, and low values (<-0.5) are cooler, La Niña conditions (NOAA, 2023). We downloaded ONI data using the data retrieval package, ‘rsoi’ (Albers 2023; version 0.5.5).

For CCE conditions, we used the Bakun Upwelling Index and SST. The Bakun Upwelling Index calculates Ekman transport to estimate upwelling intensity, with positive values implying upwelling and negative values indicating downwelling (Bakun 1973). We summarized the Bakun Upwelling Index into regions: North of CCE, Northern CCE, Central CCE, Southern CCE, and S. Baja. SST provides information on upwelling within a region, which affects ocean productivity and can support potential seabird prey (Thayer & Sydeman 2007). We used SST (°C) data from the Hadley Centre Global Sea Ice and Sea Surface Temperature (HadISST, Rayner et al., 2003) obtained from National Center for Atmospheric Research (https://rda.ucar.edu/datasets/ds277.3). We used the 1° monthly measurements calculated from a combination of in-situ and adjusted satellite-derived measurements. We generated subsets of SST data by regions using previously mentioned shapefiles (Gulf of CA and CCE Large Marine Ecosystem). We then categorized data in the CCE region into North, Central, South, and S. Baja based on previously mentioned boundaries.

For S. Baja air temperatures, we used GHCN_CAMS Gridded 2 m Temperature (°C) data which contains monthly means of high resolution (0.5 x 0.5°) analyzed global land surface temperatures (1948-2022; Fan and van den Dool, 2008). We extracted air temperatures from S. Baja as a proxy of source conditions that may have influenced Sulids to move northward.

### Data analysis

We analyzed trends in Sulid abundance and range, and investigated relationships between their abundance and environmental conditions. We analyzed each species separately, although we combined data on Masked and Nazca due to their recent split as distinct species based on behavior, breeding, and distinct bill color; yellow in Masked and orange-pink in Nazca (Pyle 2020; Pitman and Jehl, 1998).

As all four data types had different coverage, methods of collection, effort across months and years, and protocols for approving records, we could not directly compile all data together. Instead, we standardized data to include the date, species, counts of individuals, coordinates of observation, and assigned region (e.g., north CCE), then combined all data types. We produced 50 km^2^ grid cells over our study area (Figure 1; -144°W to -109°W, 21°N to 60°N) and assigned Sulid data to the corresponding grid the observation location intersected with using the ‘sf’ package (Pebesma 2018). To limit the same individuals from being double-counted in the data compilation, we used 50 km, as typical Sulid foraging ranges are 40-100 km (Lerma et al., 2020; Mendez et al, 2017; Weimerskirch et al., 2008). We then used the maximum number of Sulids seen at one time within each grid cell for each month and year of the compiled data as a metric for Sulid abundance. Although true “absence” within a grid cell cannot be obtained, we gave each grid cell a zero count during a month and year where there was any effort within the grid cell, but no Sulids detected. Lastly, SST data were extracted at each grid cell midpoint, we connected each grid cell to the nearest latitude for Bakun Upwelling Index, and remaining environmental data (ONI, S. Baja and Gulf of California SST and S. Baja air temperature) were assigned to all grid cells.

We used the gridded data within only the CCE in our analysis, which included 533 grid cells. Because there were differences in years covered by each data type (*eBird* 1935-2022, RBC 1935-2022, SEFI 1967-2022, At-sea 1980-2022), we could not analyze changes over time across the entire period. Therefore, we only analyzed years since *eBird* began (2002) and with overlap among all data types (2002-2022), which resulted in 9,712 individuals.

Prior to analysis with the environment, we evaluated the correlation between Sulids and lagged source condition. If Sulids respond to conditions in S. Baja or the Gulf of California, it may take time for them to decide to travel north, therefore, we tested relationships between their abundance with 1-12 month lagged source data (SST and air temperature in S. Baja and SST in the Gulf of California) using the Spearman rank correlation (r) calculated with the ‘cor’ function (R Core Team 2020).

We used zero-inflated Poisson generalized additive models (GAM) to investigate non-linear relationships between Sulid abundance and environmental conditions. We conducted GAMs using Restricted Maximum Likelihood (REML) smoothness estimation using the ‘mgcv’ package (Woods 2017). To determine the model of best fit, we first ran models with only the two highest correlated lagged variables (potential source conditions) and selected the month lag that best explained the variability in the data. As Red-footed, Masked, and Nazca are not found in the Gulf of California, we did not use Gulf of California SST in our analyses with these species. We checked for collinearity among all variables using variance inflation factors (VIF), which measure the amount of variance of a variable that is inflated by its correlation with another variable. We used a conservative threshold, with values less than 3 indicating no strong collinearity, and excluded variables with values greater than 3 from the overall model (Zuur, Saveliev, & Ieno 2012; Johnston et al., 2018). There was high collinearity between S. Baja SST, Gulf of California SST, and S. Baja air temperatures, therefore, we ran full models (with all other variables) for each of these variables separately. We then removed any non-significant variables, and compared each to the initial models. Variables with effective degrees of freedom (EDF; i.e., the non-linearity of a curve) equal to one were incorporated without smoothers. To select the model of best fit, we compared models using Akaike information criterion (AIC; Akaike 1973; Venables and Ripley 2002).

To test our hypothesis that Sulids increased with warm water periods, we investigated differences in Sulid abundance between years prior to extreme MHWs (2002-2012) to years during and after MHWs (2013-2022). MHWs are prolonged periods of anomalously warm waters (Hobday et al., 2016) mostly due to atmospheric forcing that affect wind and pressure systems that reduce upwelling, therefore stratifying the water column and leading to overall warming (Di Lorenzo & Mantua 2016). We compared the mean and max abundance of each species between the pre and during/post MHW periods. We calculated the percentage difference between the means and conducted a student’s t-test to compare them.

### Range shift analysis

To test whether the ranges of Sulids had significantly expanded, we first calculated the maximum latitude of 50 km^2^ gridded sightings of each species per year. We then conducted linear regression between the maximum latitudes of a species observation within 50 km^2^ grid cells per year. By consolidating observations into grid cells in this analysis, we limited the influence of differences in effort and accounted for potential double counts of the same individual.

We tested for changes in range area by counting the number of 50 km^2^ grids per year in which at least one individual was reported for each species. We controlled for effort by dividing the number of grids with an observation by the number of grids that had any effort. We used a generalized linear model with a gamma, log-linked distribution to evaluate changes overtime in range area for each species. We also compared the number of grids each species was observed within the years before (2002-2012) to years during and after MHWs (2013-2022). We calculated the percentage difference between the means and conducted a student’s t-test to compare them.

## RESULTS

### Changes in abundance

All Sulids in this study increased in abundance in the CCE (2002-2022; Figure 2). Prior to 2002, Sulids were rare vagrants north of Mexico, with only periodic observations, including a pulse of Blue-footed (mostly inland) in 1973 (Figure 2). After 2002, Cocos increased throughout biogeographic regions (from 0.21 mean number of birds per grid cell prior to 2002 to 3.80 mean birds between 2002-2022), with the highest abundance in 2014 and 2015 (5.36 mean number of birds during these two years). Since 2002, Blue-footed were infrequently observed, however, were present in periodically high abundances (e.g., 850 total birds reported in 2013 compared to 103 before 2013). It was not until 2018 that Red-footed increased from 165 total reported birds pre-2018 to 586 birds post-2018, Masked from 194 to 391 and Nazca from 135 to 634.

**Figure 2:**
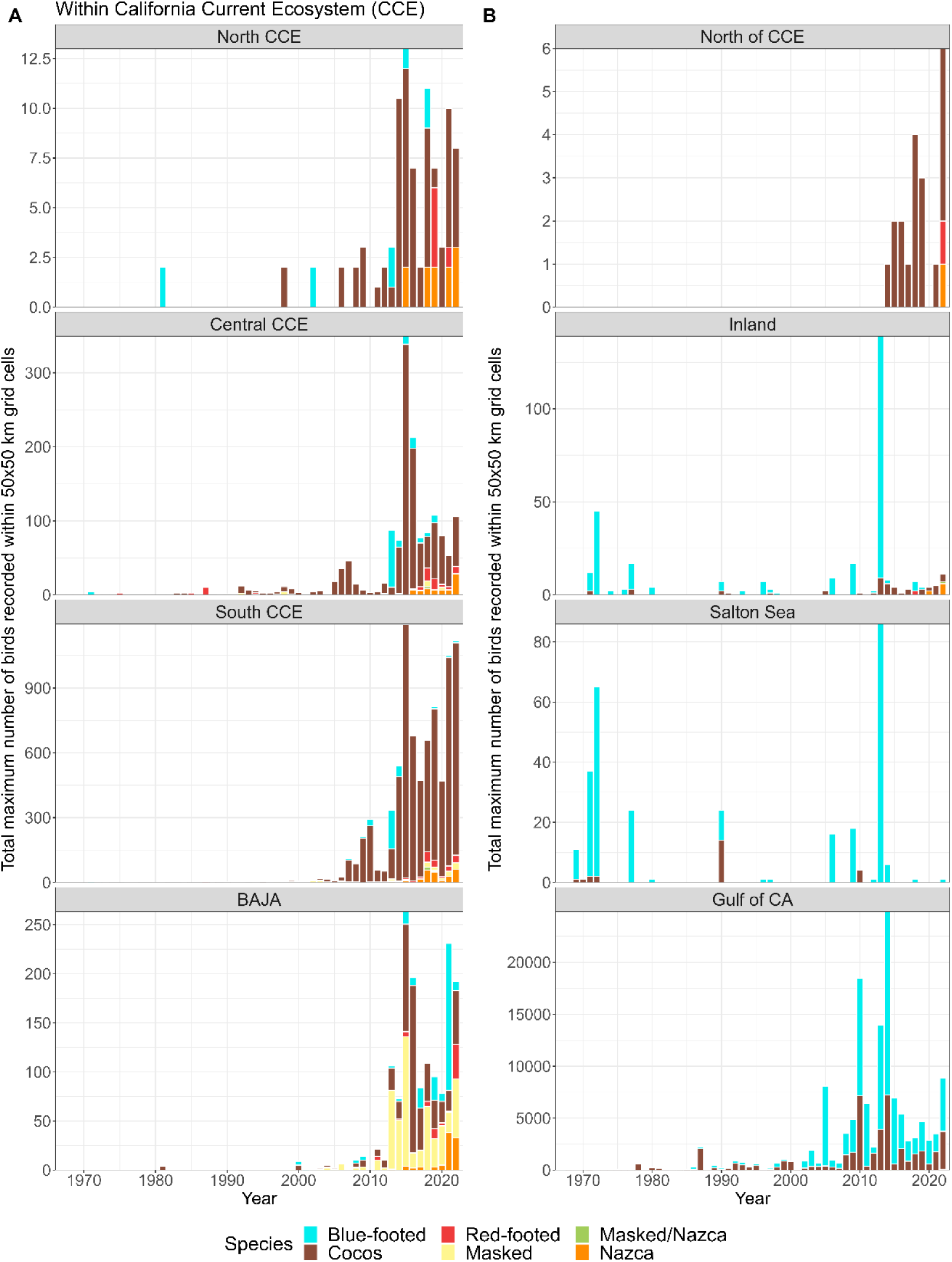
The sum of the maximum number of each Sulid recorded within a 50 km^2^ grid cell for each year and region using most of the available data (1966-2022; excluded <1966 due to low observations). (A) Regions within the California Current Ecosystem (CCE) and (B) regions outside of the CCE.

There were differences in the seasonality of Sulid occurrence. For Cocos and Blue-footed, there was no seasonality in their occurrences in the CCE, however they were more abundant north of CCE, inland, and at the Salton Sea during summer and fall, while they were more abundant between fall to spring in the Gulf of California (Figure 3S). However, Red-footed, Masked, and Nazca occurred in the CCE and S Baja mostly during late spring through fall (June to November).

**Figure 3:**
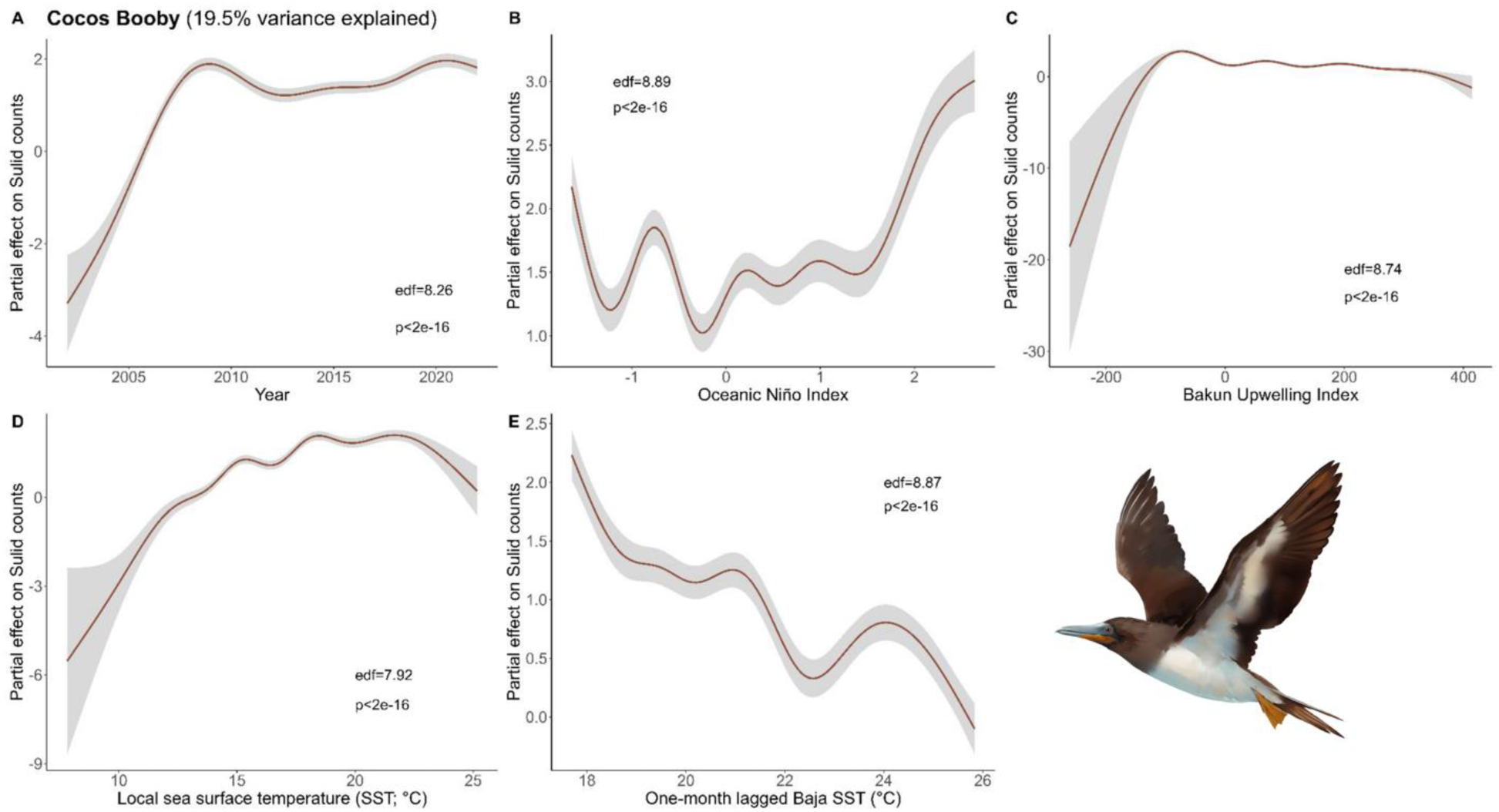
Results of generalized additive models (GAM) on Cocos Booby abundance across (A) years and with environmental variables: (B) Oceanic Niño Index, (C) Bakun Upwelling Index, (D) the sea surface temperature (SST) of the coastal latitude of each observation, and (E) one-month lagged SST in southern Baja. We used all data types to derive the maximum Cocos Booby counts within a 50 km^2^ grid cell and used these values as conservative estimates of bird abundance in our analysis (Bird illustrations by Freya Hammar).

Conservatively, using the maximum number of each species within a 50 km^2^ grid cell, all five species increased in abundance (Figure 3A-6A). The increase in Cocos was most apparent from 2002 to 2010 (Figure 3A). A similar increase occurred in Blue-footed through this same period (2005-2010; Figure 4A). Red-footed had low abundance, with high uncertainty in their estimates from 2002-2013, representing the sparse observations during this period (Figure 5A). In 2014, Red-footed abundance increased until 2021. Masked and Nazca declined from a few records in 2002 to minima in 2006 before increasing throughout the rest of the survey period (Figure 6A).

**Figure 4:**
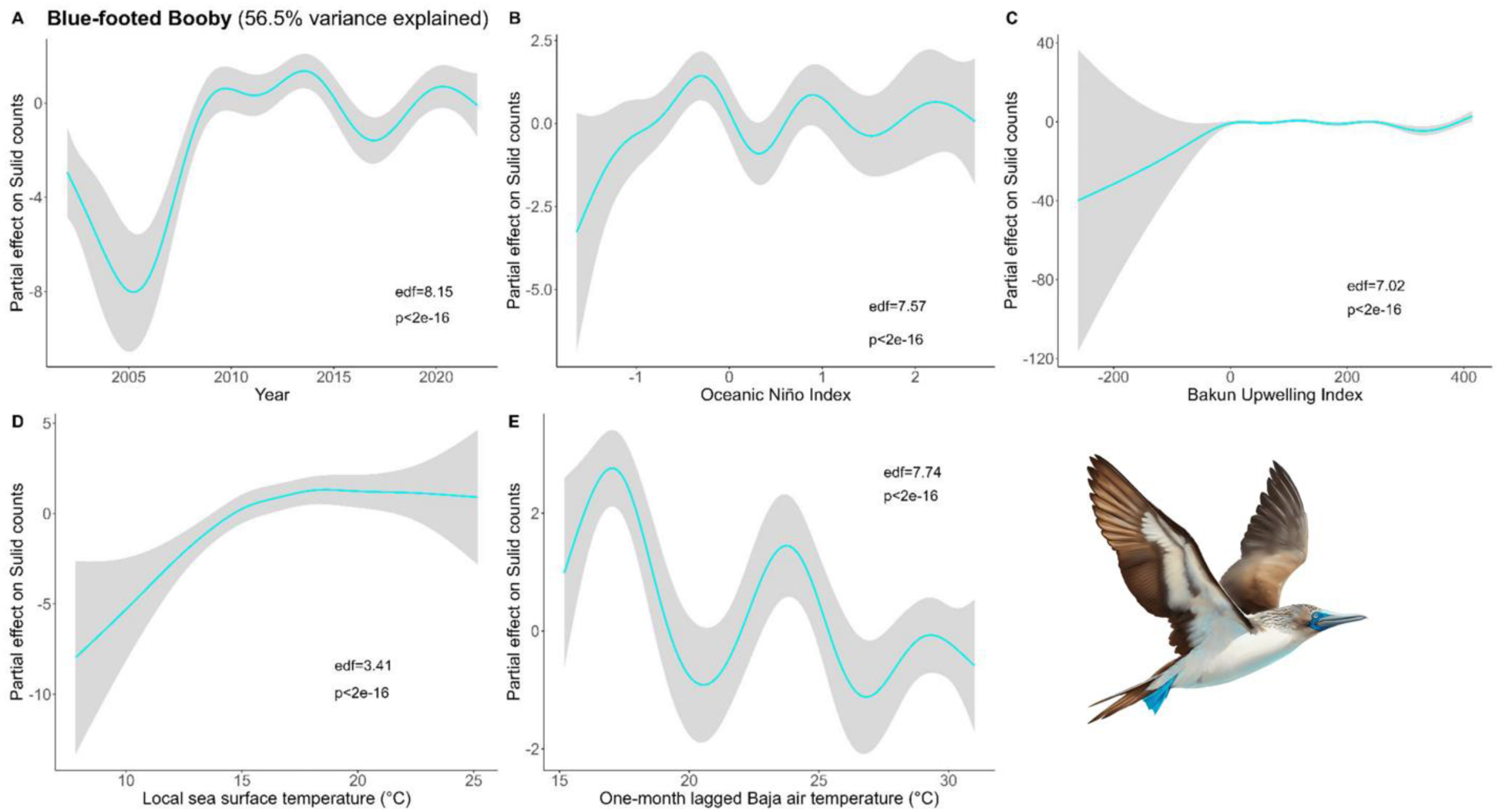
Results of generalized additive models (GAM) on Blue-Footed Booby abundance across (A) years and with environmental variables: (B) Oceanic Niño Index, (C) the sea surface temperature of the coastal latitude of each observation, and (D) one-month lagged air temperature in southern Baja. We used all data types to derive the maximum Blue-footed Booby counts within a 50 km^2^ grid cell and used these values as conservative estimates of bird abundance in our analysis (Bird illustrations by Freya Hammar).

**Figure 5:**
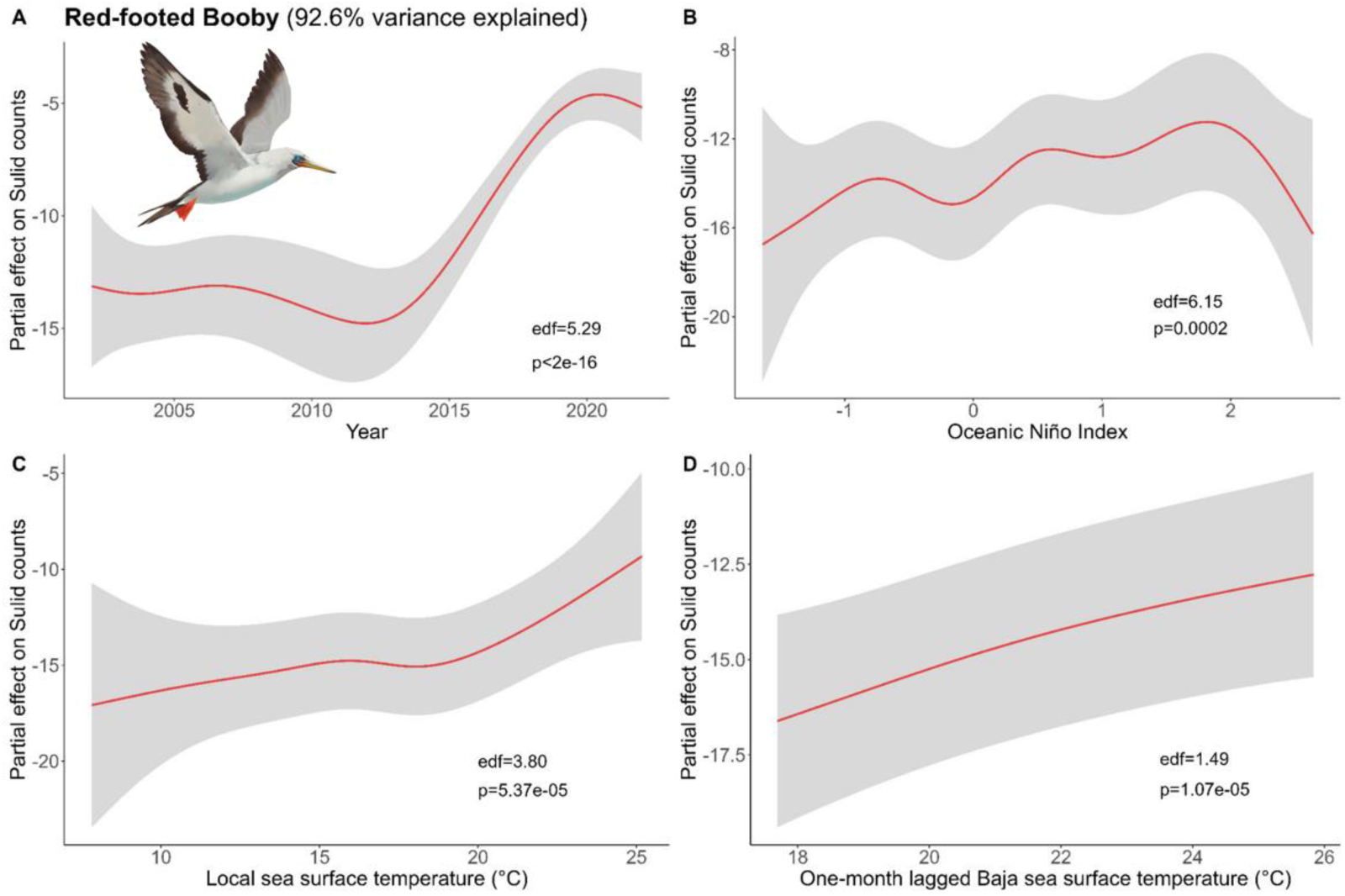
Results of generalized additive models (GAM) on Red-Footed Booby abundance across (A) years and with environmental variables: (B) Oceanic Niño Index, (C) the sea surface temperature (SST) of the coastal latitude of each observation, and (D) one-month lagged southern Baja SST. We used all data types to derive the maximum Red-footed Booby counts within a 50 km^2^ grid cell and used these values as conservative estimates of bird abundance in our analysis (Bird illustrations by Freya Hammar).

**Figure 6:**
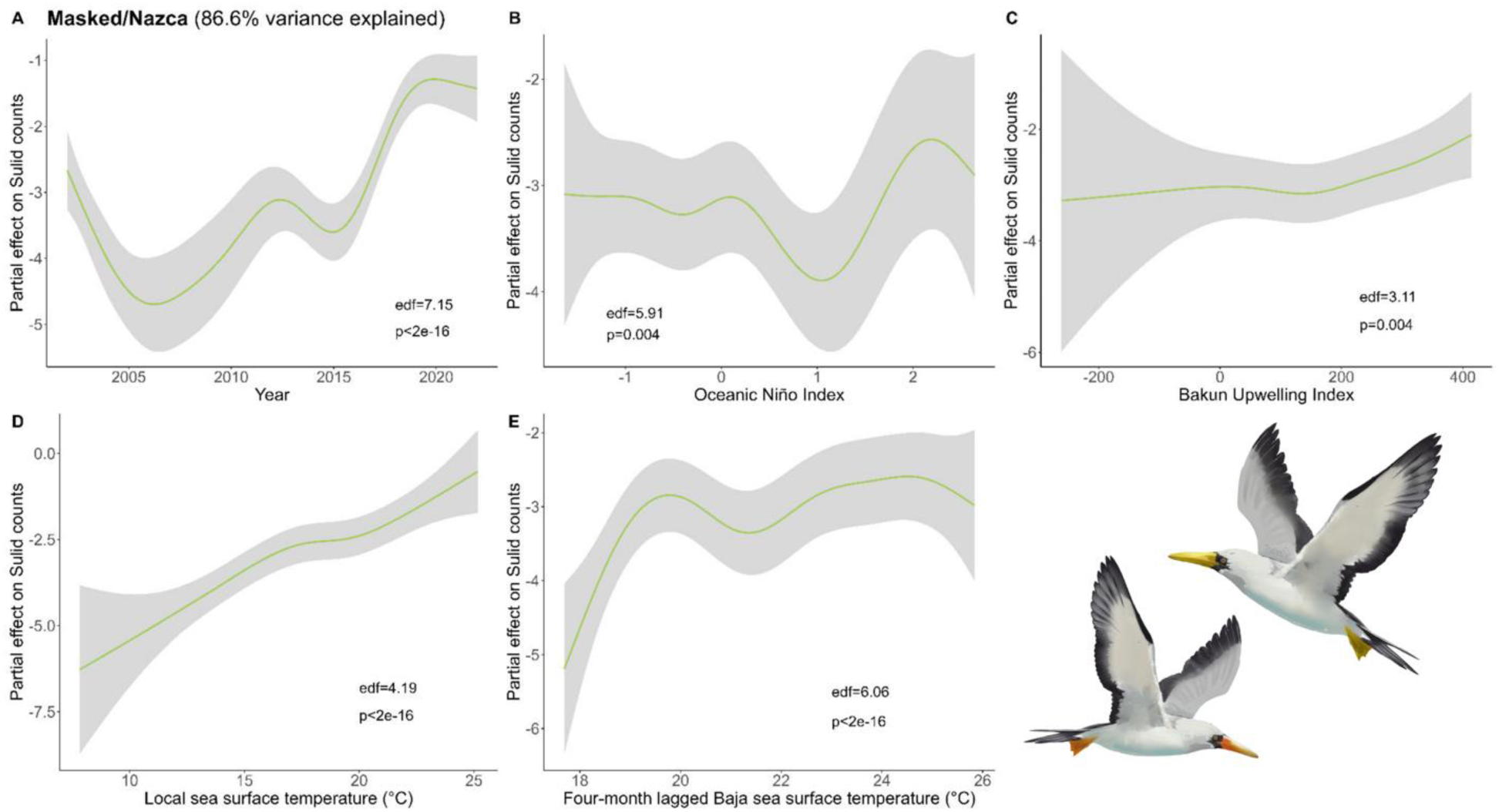
Results of generalized additive models (GAM) on Masked and Nazca Booby abundance across (A) years and with environmental variables: (B) Oceanic Niño Index, (C) Bakun Upwelling Index, (D) the sea surface temperature (SST) of the coastal latitude of each observation, and (E) four-month lagged SST in southern Baja. We used all data types to derive the maximum Masked and Nazca Booby counts within a 50 km^2^ grid cell and used these values as conservative estimates of bird abundance in our analysis (Bird illustrations by Freya Hammar).

### Changes with environmental conditions

We selected lagged variables that had the two highest correlations with the abundance of each species (Figure S4). Abundance had a positive relationship with recent conditions (0-1 month lags) and conditions 5-7 months prior for all species. We selected lagged variables with a correlation coefficient over 0.1; no correlations were greater than 0.5.

For Cocos, the final model included the variables Year, ONI, local SST, local Bakun Upwelling Index, and one-month lagged S. Baja SST (Figure 3). Our models did a poor job capturing their abundance in the CCE (19.5% variance explained), which suggests there were other factors driving this northward expansion or that their sustained presence and abundance over this time may have masked trends.

For Blue-footed, the final model included Year, ONI, local Bakun Upwelling Index, local SST, and one-month lagged S. Baja air temperature (Figure 4). Blue-footed were abundant during diverse conditions in the CCE. Local SST had the largest partial effect, with greater abundance during warmer local SST. There were pulses of Blue-footed during cool, moderate, and warm S. Baja air temperatures, indicating this is not a main driver for birds in the CCE.

For Red-footed, conditions with one-month lag had better explanatory power for their abundance than longer lagged data. The Bakun Upwelling Index was not a significant predictor and was removed, which improved the explanatory value of our model, which included Year, ONI, local SST, and one-month lagged S. Baja SST (Figure 5). Red-footed abundance was best explained by the year (increase), then local and lagged S. Baja SST. The trend with SST and ONI indicated there was higher abundance during warmer conditions (higher SST and El Niño).

For Masked and Nazca, longer lagged source data were better explanatory variables (seven-month lagged S. Baja air temperature and four-month S. Baja SST; Figure 6). The four-month lagged S. Baja SST resulted in the best fitting model, and our final model included year, ONI, local Bakun, local SST, and four-month lagged S. Baja SST. Masked and Nazca abundances were greatest during warm water conditions (local and four-month lagged S. Baja SST) and El Niño.

When we compared Sulid abundance before and after the 2013-2016 MHW, we found that all species significantly increased after this warm-water event (692-3015%; Table 1). For all species, both the mean number of individuals, and number of positive observations (lower proportion of zeros) increased (Table 1). For all species except Blue-footed, there was also a significant increase in maximum number of birds seen at one time (Cocos from 20.5±18.2 to 136.9±46.2; Red-footed from 1±0.5 to 3±0.9; Masked and Nazca from 2±0.5 to 10±2.6).

**Table 1:** Comparison between Sulid (Booby) observations before the 2013-2016 marine heatwave (2002-2012, n= 4,581grid cells) and the period during and after the heatwave (2013-2022, n=4,100 grid cells). Included are the proportion of zero observations per survey record as a measure of absence, the mean counts per period, the percentage change between the two periods, and t-test results from a comparison between these means.

| Species | Period | % Zeros | Mean counts [95% CI] | % change | t test |
| --- | --- | --- | --- | --- | --- |
| Cocos | 2002 to 2012 | 0.950 | 0.210 [0.146, 0.273] | 777.0% | $t_{(4375)} = -9.10$,<br>$p < 2.2e-16$ |
|  | 2013 to 2022 | 0.730 | 1.838 [1.492, 2.183] |  |  |
| Blue-footed | 2002 to 2012 | 0.994 | 0.013 [0.003, 0.023] | 691.6% | $t_{(5443)} = -6.46$,<br>$p = 1.17e-10$ |
|  | 2013 to 2022 | 0.952 | 0.102 [0.077, 0.127] |  |  |
| Red-footed | 2002 to 2012 | 0.998 | 0.002 [0.0005, 0.003] | 3014.5% | $t_{(4330)} = -14.13$,<br>$p < 2.2e-16$ |
|  | 2013 to 2022 | 0.947 | 0.054 [0.047, 0.062] |  |  |
| Masked/ Nazca | 2002 to 2012 | 0.993 | 0.008 [0.005, 0.011] | 1405.3% | $t_{(4450)} = -16.26$,<br>$p < 2.2e-16$ |
|  | 2013 to 2022 | 0.904 | 0.118 [0.105, 0.131] |  |  |

### Changes in range

All Sulids expanded their range northward during our study period (Figure 7 & 8). As we are only evaluating these spatial changes at the northern edge of their range, we cannot say whether it is a range shift or expansion, but for the sake of brevity, we call it a northward expansion. The range of Cocos, Red-footed, and Masked/Nazca all increased in maximum latitude (Adj. R^2^ = 0.58 p=0.00004, Adj. R^2^ = 0.46 p=0.0045, and Adj. R^2^ = 0.62 p=0.00004, respectively) and range area (F_(1,19)_=52.98 p=6.62e-07, F_(1,12)_= 32.08 p=0.000, and F_(1,17)_= 29.58 p=4.42e-05, respectively). This change was not significant for Blue-footed in either their maximum latitude or area of their range.

**Figure 7:**
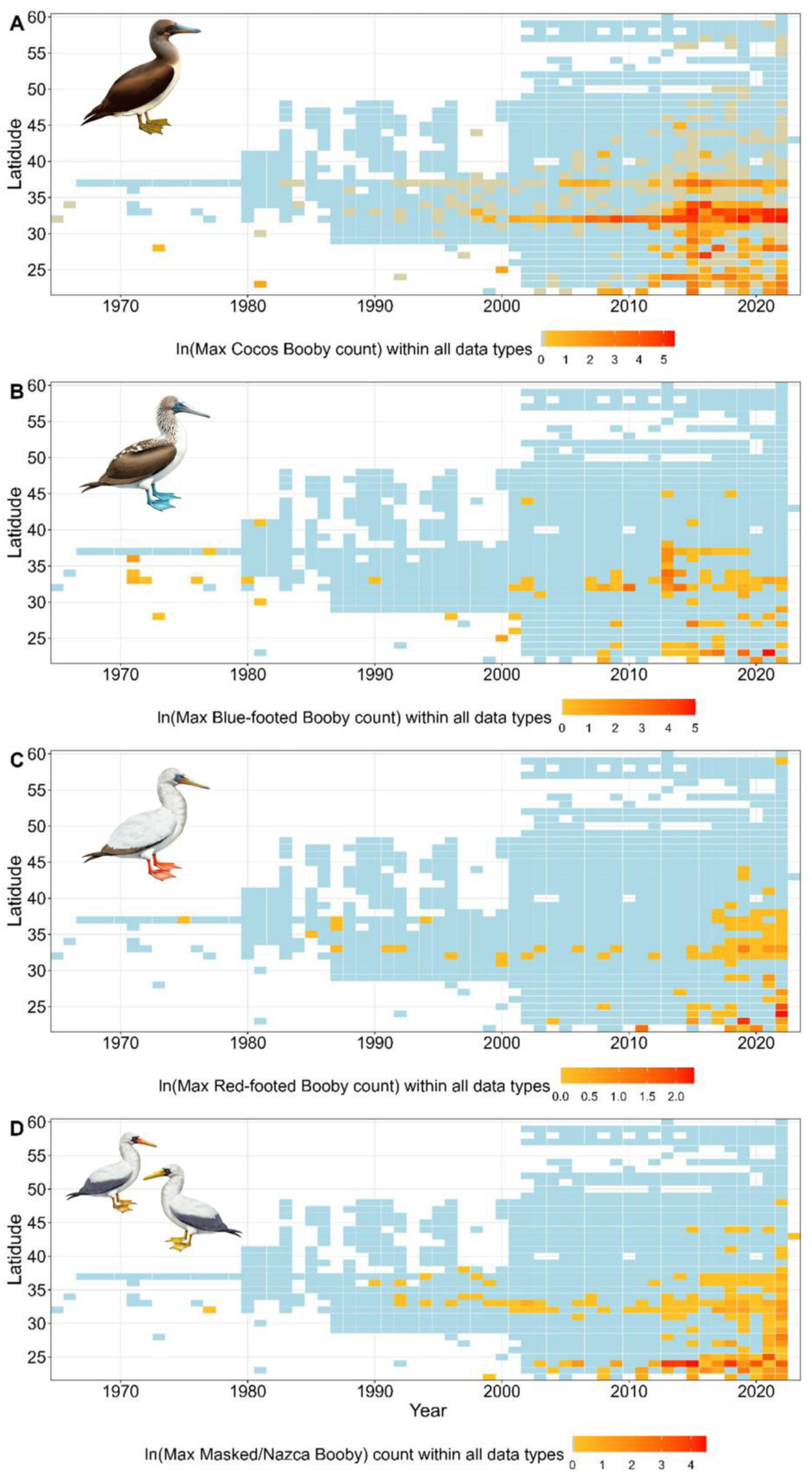
The maximum number of (A) Cocos, (B) Blue-footed, (C) Red-footed, and (D) Masked/Nazca Booby, observed within each latitude per year in the Northeastern Pacific Ocean. Boxes in blue indicate there was data collected within this latitude and year, but there were no Sulids observed. As there are many single observations and some large values, we took the log of maximum counts and centered the color gradient on the mean to show smaller values (Bird illustrations by Freya Hammar).

The northward expansion of Sulids corresponded with MHWs, with an increase in their maximum latitude by 4-6 degrees and the size of their range area by 167-777% between pre-MHW (2002-2012) and during/after MHW (2013-2022; Figure 8B & S5, Table S2). The northward expansion of Cocos began in the late 2000’s (Figure 7 & 8); there were further expansions in their range in 2015 and 2019. During the 2013-2016 MHW, Cocos abundance and maximum latitude increased (Figure 8B & S5; Table S2) and have remained at similar abundances since. Between the pre and during/post MHW periods, Cocos expanded their range area by 287.5% and the maximum latitude by 6.18 degrees. Prior to 2002, Blue-footed periodically moved northward in large pulses (Figure 7 & S5). Despite these events, there was not a significant increase in the range of Blue-footed across 2002-2022, although there was an expansion of their range (5.5 degrees and 235% area increases) between the MHW periods (Figure 8, S5). Very few Red-footed were observed within the CCE prior to 2002 (Figure 7); since then, their abundance and range have increased, especially since 2019 (Figure 8 & S5). Between MHW periods, Red-footed increased their range by 920.8% and their annual maximum latitude by 4.86 degrees (Figure 8A and Table S2). Observations of Masked and Nazca occurred sporadically throughout the 1990’s and 2000’s; their abundance increased after 2012 (Figure 7). There was an increase in their abundance during the 2013-2016 MHW; this increased further in 2019 (Figure S5). Between MHW periods, Masked and Nazca increased their range area by 1013.9% and their annual maximum latitude by 6.85 degrees (Figure 8B; Table S2).

**Figure 8:**
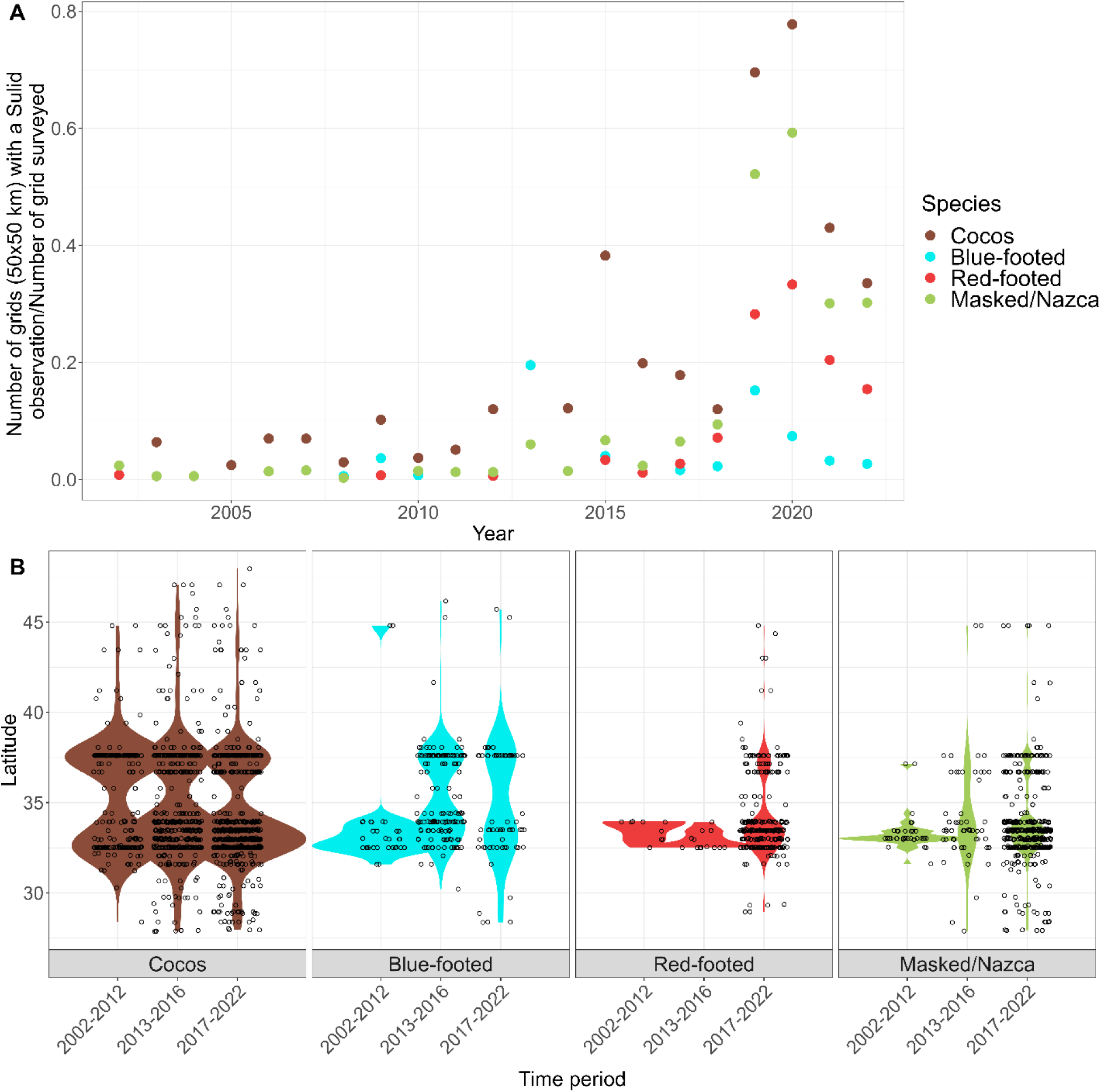
(A) The number of grid cells that a Sulid was observed divided by the total number of grid cells that had any survey effort within the CCE (North, Central and South only). In-plot statistics are the results of generalized linear models on the grids observed/grids surveyed per year for each species (COBO= Cocos, BFBO=Blue-footed, RFBO=Red-footed, and MA/NA= Masked/Nazca Booby). (B) Violin plot with overlaid observations for the latitude of sighting per month and year for each species within each period (2002-2012, 2013-2016, 2017-2022); only observations along the west coast were included (southern Baja, South, Central, and North biogeographic regions).

## DISCUSSION

Sulids increased in abundance, range area, and maximum latitude in the CCE during our study period (2002-2022), and increases in their abundance corresponded to warm water events. After increases in their abundance and range, for all species except Blue-footed, they did not contract back to their previous range.

### Changes in abundance

Cocos have an eastern Pacific distribution and were occasionally recorded along the U.S. west coast before our survey period (2002-2022). Prior to their breeding range expansion, the nearest colonies were on Clipperton Atoll and Islas Revillagigedo (Schreiber and Norton 2020) off central Mexico and on Roca Consag and Isla San Jorge in the Gulf of California (Mellink 2000). The first record in California was in 1946 (McMurry 1948) in Imperial County and since then they have shown up in small numbers until the 2000’s (McCaskie 1970). Since 2002, their abundance increased, with substantial increases after 2013 (Figure 2). This is now the most common Sulid in the CCE, with no apparent seasonality in their occurrence; they are now permanent, year-round residents.

Blue-footed are found within the Eastern Tropical Pacific and have a coastal distribution compared to the other Sulids in our study (Nelson 1978). Prior to their breeding range expansion, the closest breeding locations were within the Gulf of California, although the largest breeding colonies occur on the Galapagos Islands (Nelson 1978; Díaz and Gómez 2020; Howard et al., 2024). This species was first recorded in California at the Salton Sea in 1929 (Clary 1930) and is the most commonly observed species inland. Although there were large pulses in their abundance in 1973 (inland only) and in 2013 (all regions but the north), their distribution and abundance trends are different from other Sulids as they have not significantly increased overtime or remained in large numbers after warm water events. As they are a coastal Sulid, there may be limitations on their range and/or specific habitat preferences responsible for this difference from other Sulids.

Both Red-footed and Masked have a pantropical distribution, while Nazca are only found in the Eastern Tropical Pacific and all three are highly pelagic. Their nearest breeding locations occur on Islas Revillagigedo and/or Clipperton Atoll off central Mexico (Howell and Pyle 1997; Pitman and Jehl 1998). These Sulids were mostly observed during summer and fall. Masked and Nazca are seasonal breeding species. For example, Masked that breed on Clipperton Island begin laying eggs in November and chick-rearing can continue through February (Weimerskirch et al., 2008). With Nazca that breed on the Galapagos, their breeding season spans from October to June (Tompkins & Anderson 2021). Therefore, most of the birds showing up in the CCE are arriving (May-Dec) during the non-breeding season. Conversely, Red-footed are seasonal breeders at some locations (Weimerskirch et al., 2005), while not at others (Nelson 1969). Their seasonal occurrence may indicate that birds in the CCE are from colonies with seasonality, and they are here during the non-breeding season. There were limited sightings of these three species during most of our study period. The first Red-footed observation in California was at SEFI in 1975 (Figure 1S, Huber and Lewis 1980), first Masked off San Clemente Island in 1977 (Lewis and Tyler 1978), and first Nazca was in 2001 on a ship in San Diego Harbor (Benson et al., 2022). Reports of these species remained sparse before increasing in the mid-2000’s, and substantially since 2018.

### Changes with environmental conditions

As seabird distributions can reflect productive conditions for their prey, we evaluated which environmental conditions may have encouraged changes in Sulid ranges. Connections between marine predators and the environment can be concurrent, or they may occur at different lagged time scales depending how the distribution, abundance, and quality of their prey are affected (Wakefield et al., 2009; Mendez et al., 2017A). We found that Sulids responded to their ‘source’ conditions (SST and air temperatures) at scales of 0-1 months or 5-7 months, such that some individuals appear to respond rapidly to changing conditions, while others appeared to wait to head north until conditions worsened.

Our models performed well in characterizing the environmental conditions that may drive the abundance of Red-footed, Masked and Nazca, however the trends for Cocos and Blue-footed were not as straightforward. Although Cocos increased during and after MHWs, trends with environmental variables were complicated by their year-round presence in recent years. Similarly, Blue-footed did not show a clear trend with the environment, although a pulse in Blue-footed abundance occurred during a warm period (2013) when conditions further south or in the Gulf of California may have been poor compared to conditions in the central and northern CCE. These more favorable conditions may have retained recurrent visitors, such as the two second-year birds showed up on SEFI in 2014 and came back together to the same location each year (2015-2019), and the new breeding pair on Sutil Rock (Howard et al., 2024).

For the highly pelagic and wide-ranging Red-footed, their abundance increased between the mid-2010’s to 2022 during the warm water and reduced upwelling conditions of the extreme MHWs. Although there was a positive relationship between Red-footed abundance and one-month lagged S. Baja SST, some of these individuals may have travelled from breeding locations in Hawaii, which may explain the one record of this species off Alaska in 2022.

Masked and Nazca abundance was higher during warm-water conditions. Tracking studies have shown that these large Sulids will travel far in search of productive conditions (Sommerfeld et al., 2015); their occurrence during poor conditions may be an indication that conditions in their typical range were even worse.

### Changes in Sulids range

There has been a northward expansion of the five Sulids in this study into the CCE. Both the range area and the maximum latitude have increased overtime, for all except Blue-footed. Although there were increased area and latitude of Blue-footed between pre/post MHW, it was not a linear trend. Blue-footed presence here is limited except for a few substantial incursions; the few individuals that remain afterwards show up as vagrants. This pattern may change now that they expanded their breeding range north; however, breeding appears to be limited to only a few nests at most (Díaz & Gómez, 2011; Howard et al., 2024). Prior to the northward breeding expansion to Sutil Rock, Cocos had expanded their breeding range to Middle Rock, Baja California in 2005 (Whitworth et al., 2007). After this initial northward breeding expansion, Cocos abundance rapidly increased, with peak abundance occurring during the initial years of attendance and breeding attempts at Sutil Rock (starting in 2013; Howard et al., 2024).

For Red-footed, their northward expansion did not occur until 2018. They were observed sporadically before 2002, as lone individuals mostly within 31-33N, with one record at SEFI in 1975 (∼37N). Starting in 2018, their range area and northward limit expanded until 2021.

Even though Masked and Nazca increased in abundance during the mid-2000’s, their range did not significantly expand northward until 2013. Initially, the expansion was due to an uptick in Nazca abundance and northern range (Figure S1 & S2). After 2018, the abundance, range area, and maximum latitude of both species increased and continued to increase to 2022. In the most recent year of our data (2022), there was at least one Masked/Nazca detected within each degree of latitude between 22°N and 37°N.

### Changes associated with marine heatwaves

There were a variety of climatic conditions during our survey period (2002-2022), including several intense MHWs. Beginning in the fall of 2013, waters in the North Pacific began to warm and it soon became the largest, longest-lasting MHW globally; reaching temperatures of three standard deviations above the mean (Bond et al., 2015; Di Lorenzo & Mantua 2016; Suryan et al., 2021; Cavole et al., 2016). Anomalous warming began in the southern CCE in the spring 2014, and by summer records off Baja California reached 4°C above the mean (Robinson, 2016; Leising et al., 2015). This MHW peaked in 2015, persisted until mid-2016, and warmer conditions remained through 2016 along the equatorial northeastern Pacific (Gentemann et al., 2017; Hobday et al., 2018).

The 2013-2016 MHW caused massive ecological changes in the North Pacific including shifts in the community composition, abundance, and/or quality of zooplankton and fishes (Brodeur et al., 2019; von Biela et al., 2019; Arimitsu et al., 2021), changes in the ranges of lower and upper trophic level organisms (Sutherland et al., 2018; Lonhart et al., 2018; Tanaka et al., 2021; Welch et al., 2023; Osborne et al., 2020), and mass mortality events of marine species (Jones et al., 2018; Piatt et al., 2020; Cavole et al., 2016). Such changes persisted in this ecosystem for at least five years (Suryan et al., 2021; Scannell et al., 2020). In addition to these impacts, there was little time for this ecosystem to recover before the next MHWs. During the summer of 2018, there was a localized MHW in the SCB that extended from Point Conception, California to Point Eugenia, Baja California Sur (Fumo et al., 2020). While this MHW was severe in the SCB, north of Point Conception SST anomalies were negative, which may have encouraged Sulids northward. The next MHW began in summer of 2019 due to a weakening of the high-pressure system in the North Pacific and persisted through 2020 (Amaya et al., 2020; Chen et al. 2021; Welch et al., 2023).

The abundance and range of all five Sulids in this study increased during and after these MHWs (Figure S5). During the 2013-2016 MHW, Cocos abundance peaked (Figure S1). This increase occurred alongside the initiation of northern breeding expansion to Sutil Rock, and the continued use of this colony supports that although Cocos had been increasing in this region before the MHW, they stayed afterwards (Howard et al., 2024). In contrast, Blue-footed increased during initial warming of the MHW (September-December 2013), however they did not remain afterwards, with the exception of a few, likely recurring individuals. Red-footed remained rare vagrants until 2018, during the severe MHW in the SCB and Baja California, when their abundance and range suddenly increased. Masked and Nazca began increasing preceding the 2013-2016 MHW, however their abundance and range substantially increased during the MHW and continued afterwards.

As MHWs are projected to increase in frequency and intensity, in addition to long-term warming, we hypothesize that these species will continue their northward expansion (Oliver et al., 2018; 2019; Laufkötter, Zscheischler & Frölicher 2020; Jacox et al., 2020). Seabird prey are poikilothermic, therefore changes in SST (along with subsequent changes in nutrients from depressed upwelling) directly influence the distribution and health of their prey (Sunday et al., 2012; Alfonso et al., 2021), and indirectly the distribution of seabirds. Changes in marine predator distributions can have consequences for pre-existing species, such as increased competition for resources, including food and breeding habitat (Pinsky et al., 2020; Petalas et al., 2021; 2024).

### Other drivers of Sulid northward range expansion

Although environmental conditions explained the trends in some species’ abundance, it did not correlate with all Sulids. Environmental variables are only proxies for conditions that may alter seabird prey; therefore, investigating prey abundance trends directly would be beneficial. As was hypothesized in Howard et al. (2024), the combination of warm water conditions and prey availability likely played a substantial role in the breeding range expansion of Cocos and Blue-footed. Coinciding with the 2013-2016 MHW was the collapse of the sardine fishery in the Gulf of California (2013-2015; Velarde et al., 2013), while the anchovy biomass around the Channel Islands was at mean conditions (Gallo et al. 2019). These factors set up prime conditions for birds in the Gulf of California to move northwest into the CCE.

In addition to oceanographic conditions and prey availability, successful conservation work may have influenced range expansions. Many of the islands with Sulid colonies within the Gulf of California and off the Pacific coast of Mexico have experienced successful eradication of invasive species (Aguirre-Muñoz et al., 2018). Invasive mammals can cause devastating harm to seabird eggs, chicks, and even adults (Spatz et al., 2023). The removals of invasive species and habitat restoration on these islands have had a positive impact on seabirds that breed there (Bedolla-Guzmán et al., 2019; Méndez Sánchez et al., 2022). An investigation into how the timing of restoration and population trends relate to Sulids trends in the CCE would be useful. An increase in population size has been shown to increase vagrancy in other seabird species (Acosta Alamo et al., 2020; Zawadzki et al., 2021), therefore, trends on these islands could play a role in their range expansion.

The potential drivers for the northward expansion of Red-footed, Masked, and Nazca are complicated by their breeding colonies being far removed from the CCE. We also do not have any indication as to what colonies these visitors are travelling from. Increased biologging of individuals at colonies in the North Pacific are needed to better determine foraging ranges during the non-breeding season and how individuals respond to environmental change.

### Reversal to the Miocene

Although warming from anthropogenic climate change is occurring at a more rapid pace, our Earth has previously experienced warm periods. During the mid-Miocene (17-15 Ma) occurred the ‘Middle Miocene Climate Optimum’ (MMCO), a period of high CO_2_ and dramatic, global warming (Foster et al., 2012; Knorr & Lohmann 2014). Fossil evidence from southern California and Baja California show that Sulids were once a dominant member of the seabird community, and had high diversity, with six species recorded in California (Warheit 1992; Stucchi et al., 2015). Sulids were abundant along the Pacific coast of South America from the early Middle Miocene, they expanded to North America during the Middle to Late Miocene (Kloess and Parham, 2017), and were present in California and Baja California until Late Miocene. Their expansion into the Northeastern Pacific occurred during the MMCO, and as temperatures decreased, the community shifted to diving species, such as alcids (Kloess and Parham, 2017). These shifts in temperature regimes and seabird communities occurred in tandem with the development of the upwelling system in the CCE (White et al., 1992; Holbourn et al., 2014). The recent influx of Sulids to the CCE during extreme warming mirrors shifts that occurred during the Miocene. As we continue to understand how human-induced increases in CO_2_ and subsequent warming affect marine ecosystems, utilizing paleo records of past events may shed light on the mechanisms and consequences of these changes.

## ACKNOWLEDGEMENTS

We thank all data collectors and contributors to this study, including at-sea data principal investigators (Table S1) and seabird observers, staff and volunteers that collected Sulid observations from SEFI, the CA Rare Birds Committee, and *eBird* reviewers. We also sincerely thank the Farallon Institute staff for the substantial data contribution from the California Cooperative Oceanic Fisheries Instigations. We extend much appreciation to the millions of birdwatching community scientists whose contributions to *eBird* make studies like this possible. We also thank the Cornell Lab of Ornithology’s *eBird* team for creating and maintaining eBird, and for making this incredible dataset freely available online, along with coding tutorials.

## SUPPLEMENTAL MATERIALS

**Table S1:**
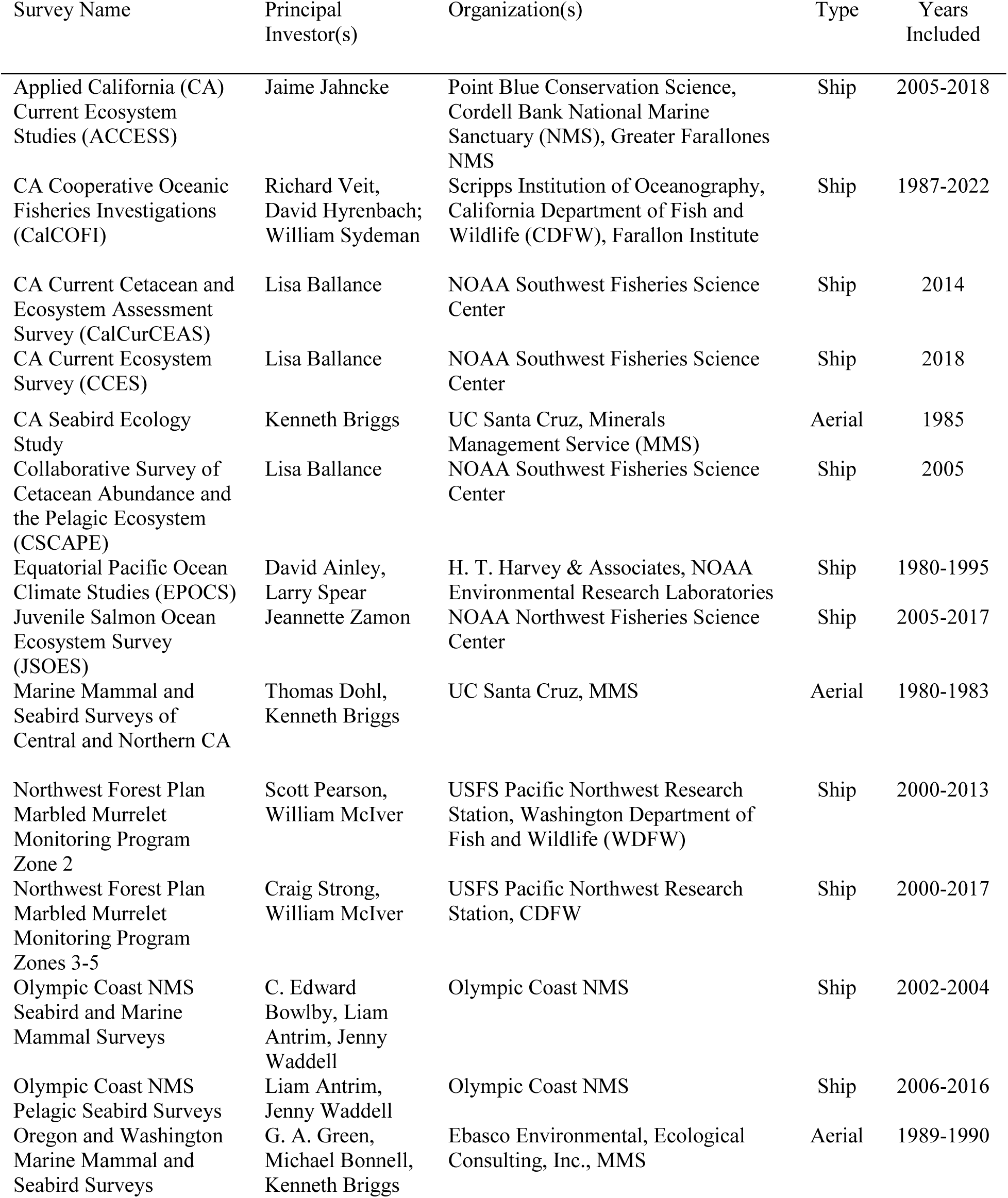
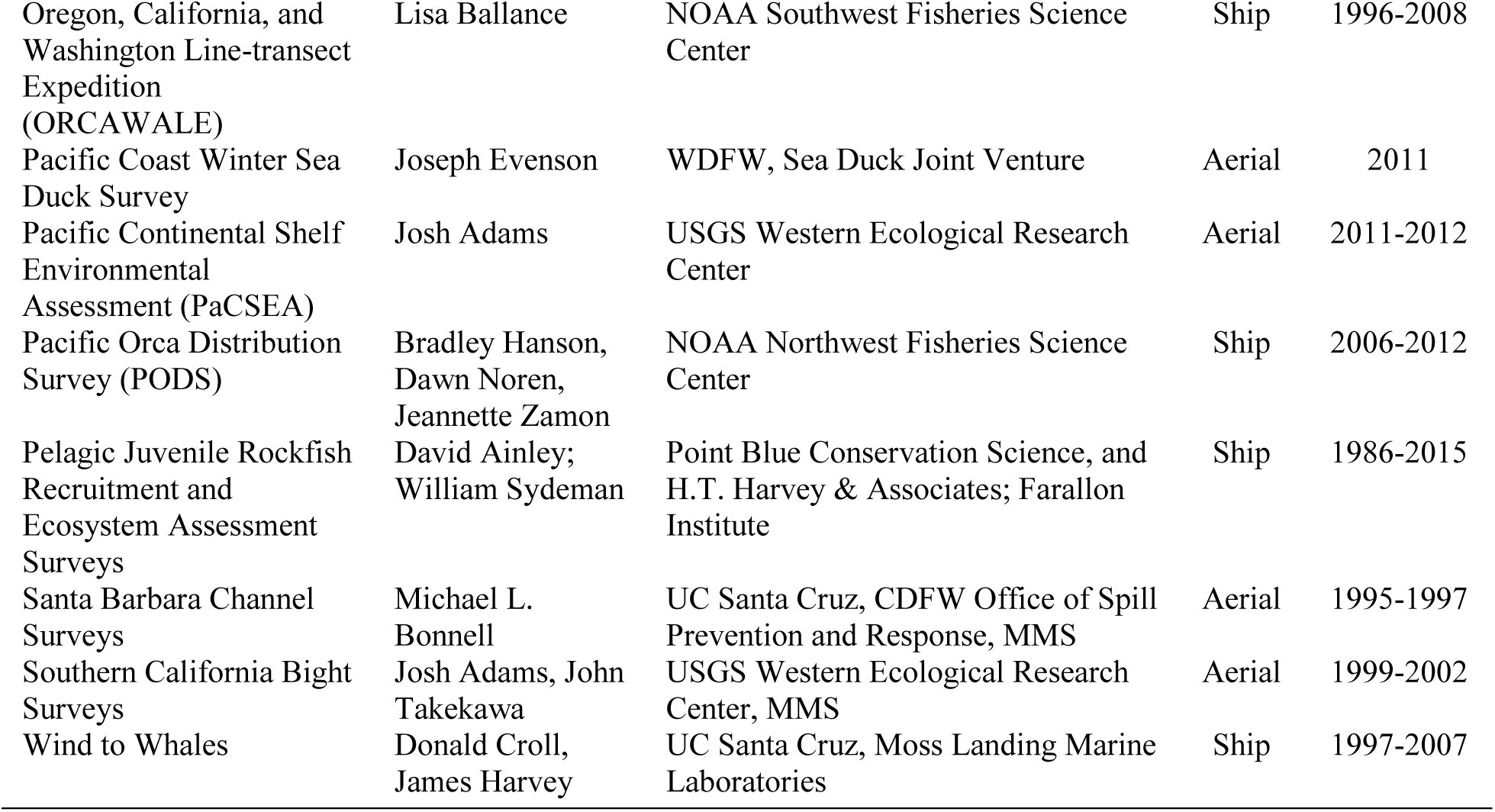
Survey Data used in this study. Much of this was compiled by Leirness et al. (2021), with added years of CalCOFI, ACCESS, and the NOAA Southwest Fisheries Science Center CCE data.

**Table S2:** Comparison between the range area of each Sulid (booby) before the large marine heatwaves (2002-2012) and the period during and after the heatwaves (2013-2022). We used the number of grids (50) a Sulid was recorded as a proxy for range area. Included are the mean annual number of grids each species was observed, the percentage change between the two periods, and t-test results from a comparison between these means.

| Species | Period | Mean # of grids [95% CI] | % change | t test |
| --- | --- | --- | --- | --- |
| Cocos | 2002 to 2012 | 9.60 [ 6.62, 12.58] | 287.5% | $t_{(12.5)} = -7.44, p = 6.26e-06$ |
|  | 2013 to 2022 | 37.20 [30.57, 43.83] |  |  |
| Blue-footed | 2002 to 2012 | 2.00 [0.76, 3.24] | 235.0% | $t_{(10.4)} = -2.04, p = 0.067$ |
|  | 2013 to 2022 | 6.70 [ 2.37, 11.03] |  |  |
| Red-footed | 2002 to 2012 | 1.20 [0.81, 1.59] | 920.8% | $t_{(7.1)} = -3.78, p = 0.007$ |
|  | 2013 to 2022 | 12.25 [6.53, 17.97] |  |  |
| Masked/ Nazca | 2002 to 2012 | 1.63 [ 1.27, 1.98] | 1013.9% | $t_{(9.0)} = -3.96, p = 0.003$ |
|  | 2013 to 2022 | 18.10 [9.95, 26.25] |  |  |

**Figure S1:**
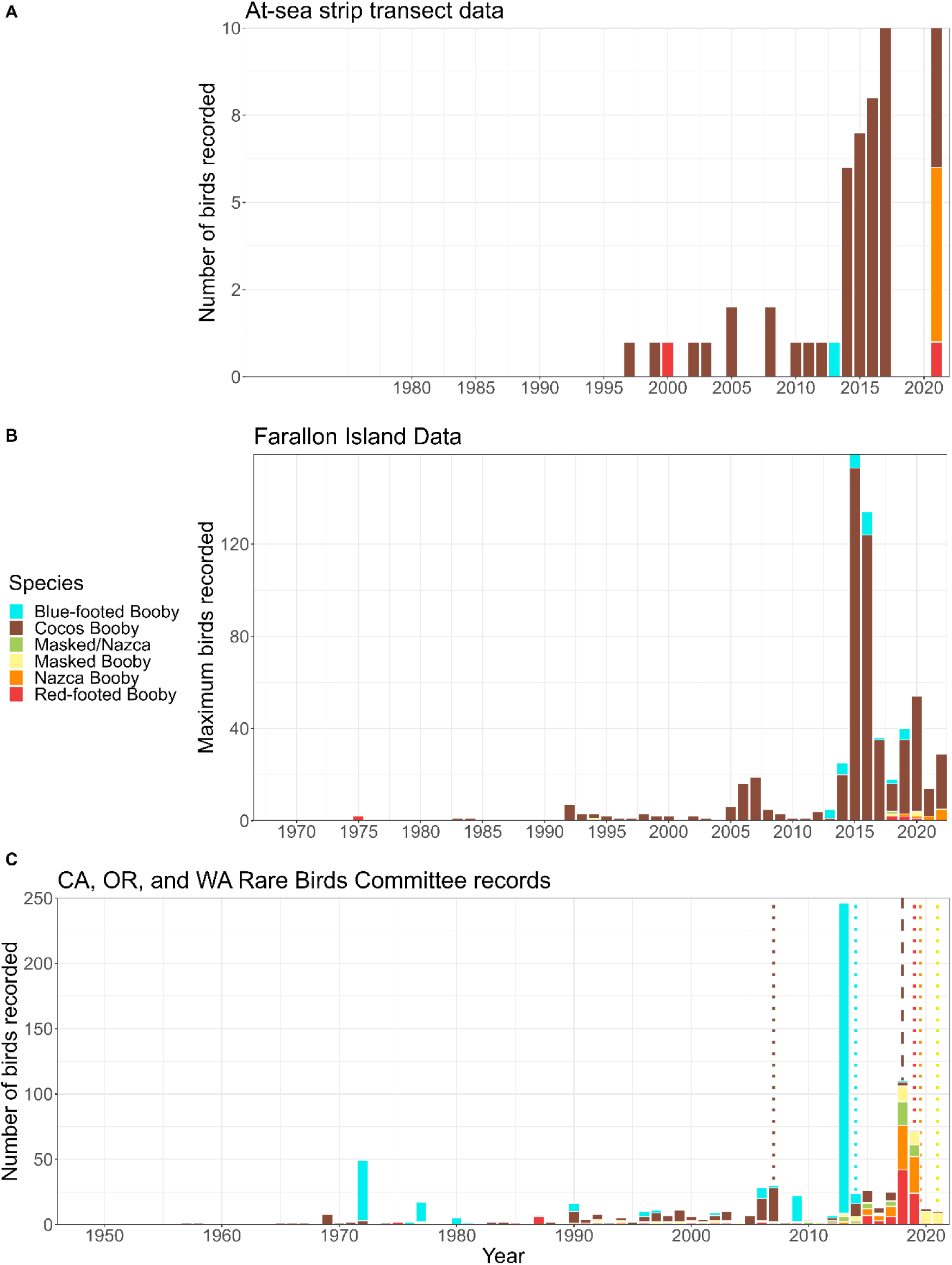
Sulid recorded in the (A) at-sea data (annual total counts), (B) the Farallon Island records (annual sum of maximum counts per month), and (C) the California, Oregon, and Washington Rare Birds Committee (RBC) records (total number of individuals). The dotted lines on the RBC plot indicate the years each corresponding colored species was removed from the California rare birds list, and the dashed line represents the year Cocos Boobies were removed from Washington rare bird lists.

**Figure S2:**
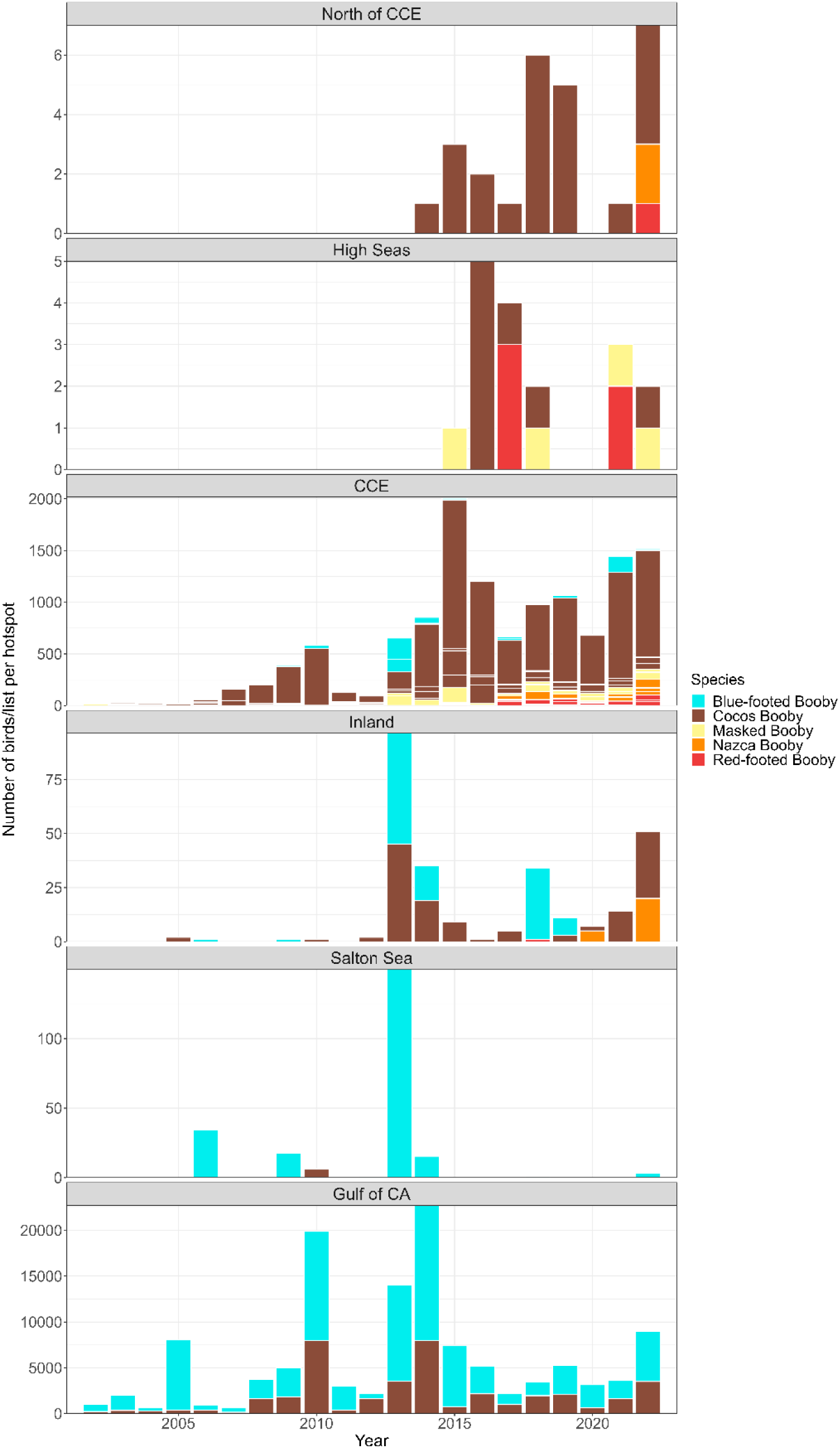
Total number of Sulid observations recorded through *eBird* (2002-2022) within the different regions within and surrounding the California Current Ecosystem.

**Figure S3:**
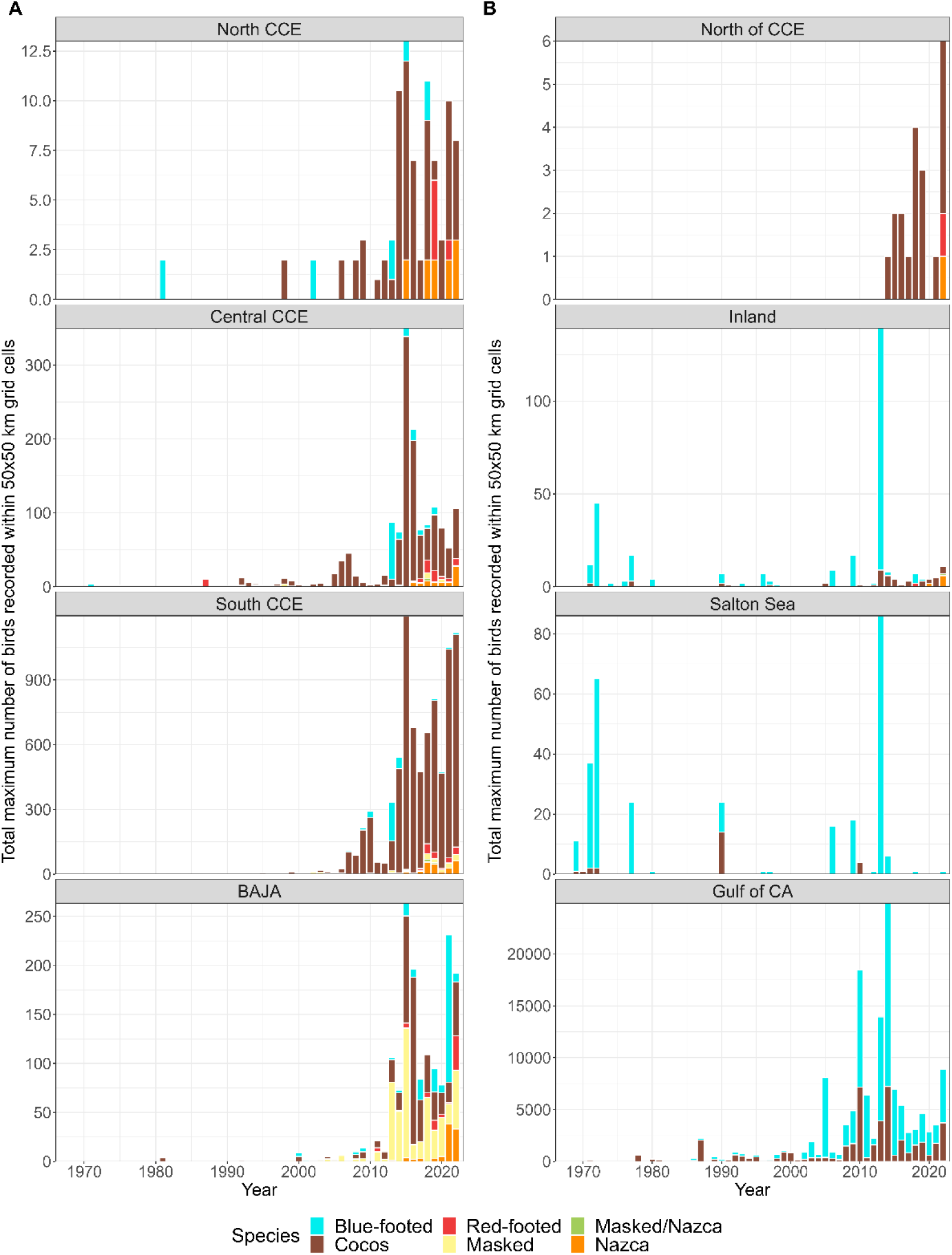
The maximum number of each Sulid recorded within a 50 km grid cell for each month and region using all available data (1935-2022). (A) Regions within the California Current Ecosystem (CCE) and (B) regions outside of the CCE.

**Figure S4:**
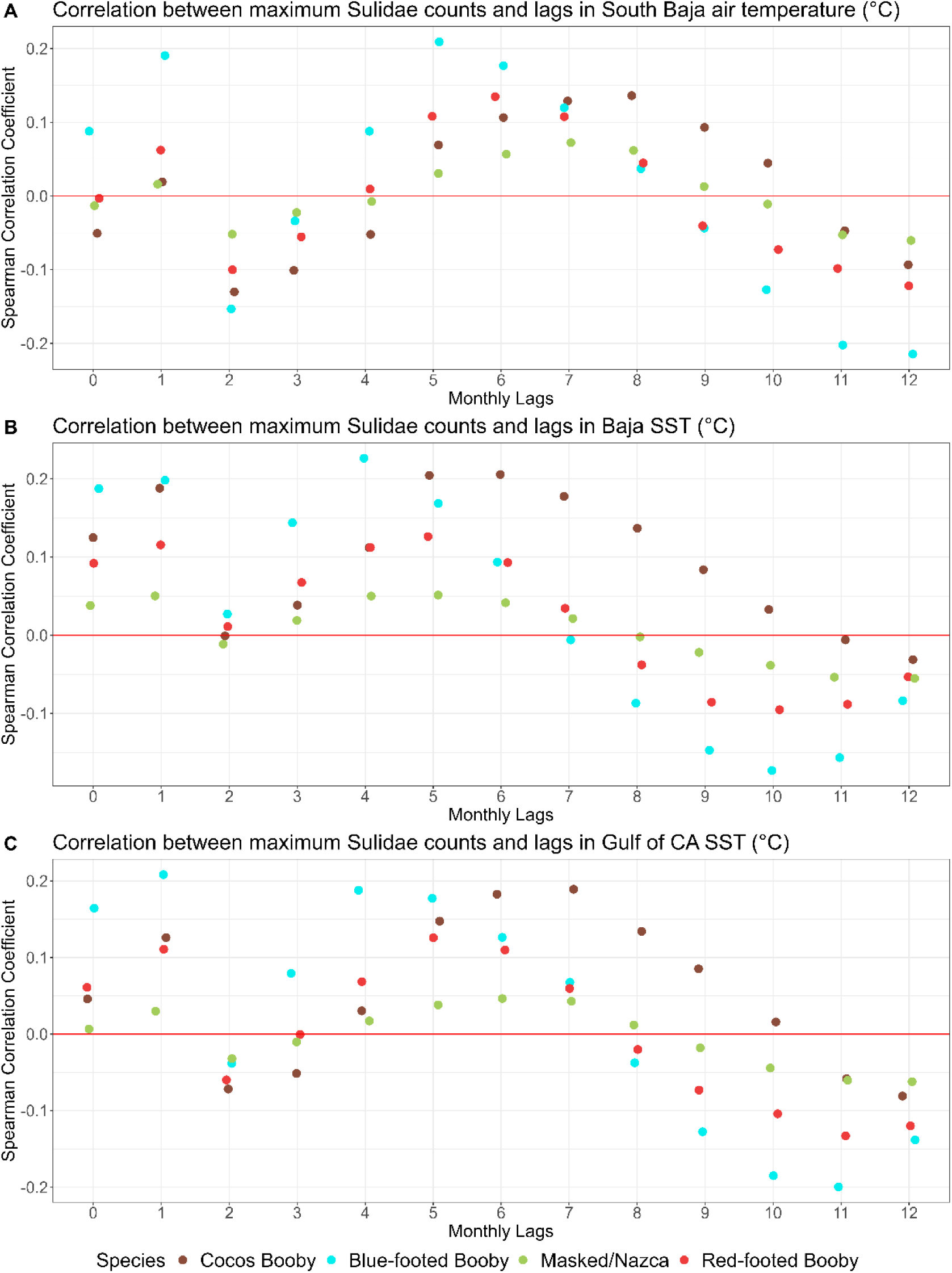
Correlations between environmental conditions around Baja California and Baja Sur, Mexico, with maximum counts of each species within the California Current Ecosystem. We compared lagged, monthly data for each variable (A) south Baja air temperature, (B) S. Baja sea surface temperature (SST), and (C) Gulf of California SST.

**Figure S5:**
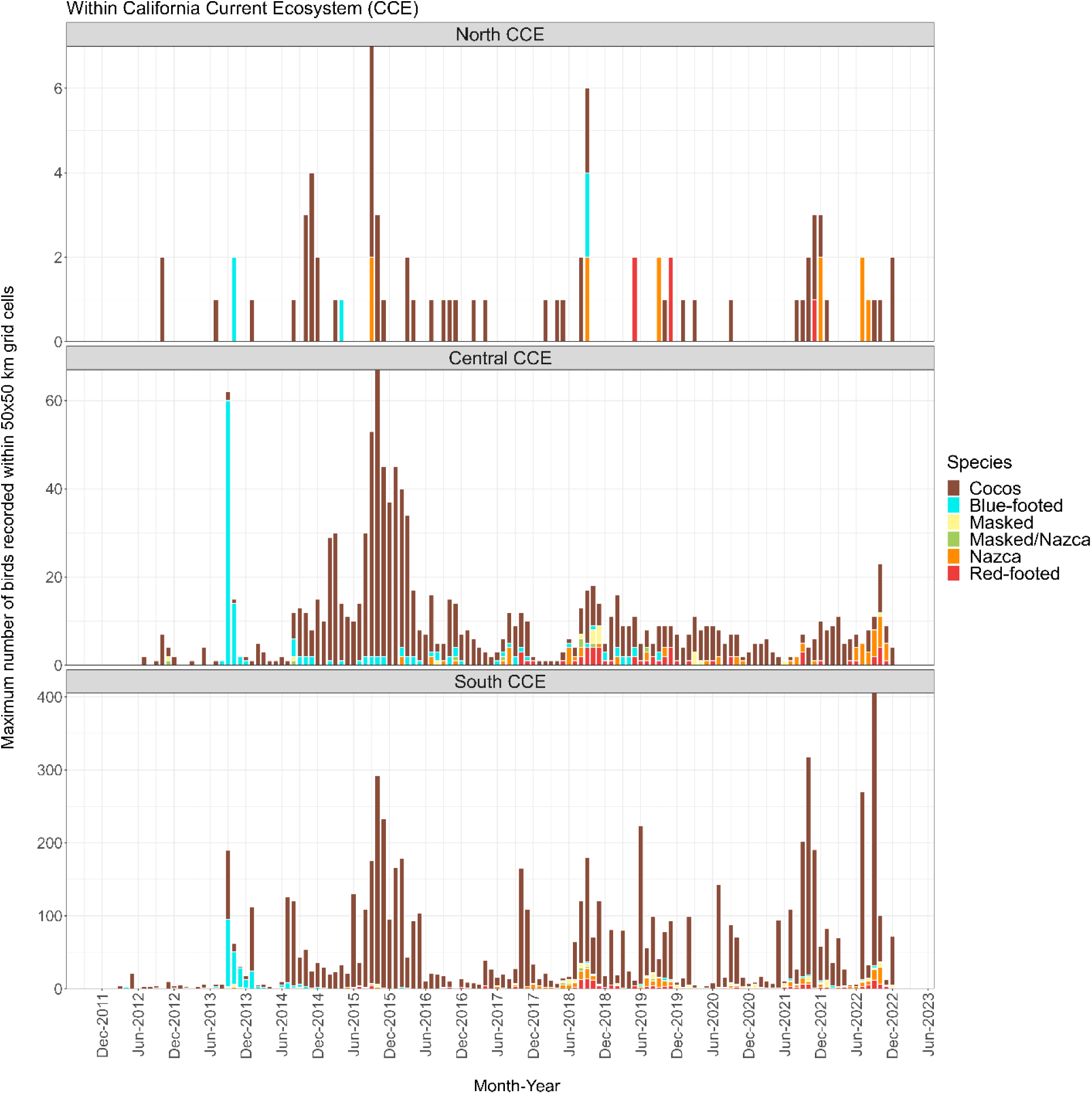
Sum of the maximum Sulid observations per month and year for each region within the CCE over the period of two marine heatwaves (2013-2016 and 2019-2020).

